# Non-apoptotic function of caspase-3 in morphogenesis of epithelial tubes of *Drosophila* renal system

**DOI:** 10.1101/2020.08.17.219634

**Authors:** Shainy Ojha, Madhu G. Tapadia

## Abstract

Cells trigger apoptosis to eliminate themselves from the system, when tissue needs to be sculptured or they detect any abnormality within them, thus preventing irreparable damage to the host. *Drosophila* Malpighian tubules express apoptotic proteins, without succumbing to cell death. Here we present evidence to show apoptosis independent role of executioner caspase, Drice, for precise architecture and function of Malpighian tubules. Drice is required for precise cytoskeleton organization and convergent extension, failing which the morphology, size, cellular number and arrangement gets affected. Acquisition of star shape of stellate cells in adult Malpighian tubules requires Drice. We demonstrate that Drice regulates expression of Rho1GTPase and localization of polarity proteins. Our study shows a probable mechanism by which Drice governs tubulogenesis via Rho1GTPase mediated coordinated organization of actin cytoskeleton and membrane stablisation. Furthermore, defective morphology of tubules leads to abnormal osmoregulation and excretory functions. Collectively our findings suggest a possible non-apoptotic function of caspase-3 in the fine tuning of cell mobility during tubule development and our results will add to the growing understanding of diverse roles of caspases during its evolution in metazoans.

## Introduction

Programmed Cell Death or apoptosis is a tightly regulated physiological process, which brings about cell death, by sequential activation of specific set cysteine proteases (Caspases), that are evolutionarily conserved across multicellular organisms [1]. During execution, the initiator caspases are activated by large multimeric apoptosome complexes, leading to the activation of effector caspases, which in turn target selected protein substrates to proteolytic degradation resulting in breakdown of cellular function and ultimately death. Execution of apoptosis in *Drosophila*, is initiated by the activation of pro-apoptotic genes, *reaper, hid* and *grim* leading to inactivation of the IAPs (Inhibitors of apoptotic proteins), which restrain the activation of caspases. As soon as the inhibition of IAPs are removed, the initiator caspase, Dronc is activated by DARK-mediated apoptosome, which then cleaves and activates effector caspases, Drice and DCP-1 culminating in irreversible cleavage of target proteins [2-4]. Since the discovery of caspases, they have been extensively studied for their role in Programmed cell death, however at variance with the existing knowledge of caspase functions, many studies now confirm that their mere presence does not qualify the cells to enter the apoptotic pathway and cells can survive despite caspase activity [5-7]. In tune with this knowledge, several studies in *Drosophila* support caspase dependent signaling in non-apoptotic processes such as, cytoskeletal remodeling and associated cellular shaping to promote major morphological changes in some sensory neurons during dendritic pruning [8, 9], cytoplasmic extrusion during spermatid terminal differentiation, thorax closure in pupal development and trachea elongation by regulating endocytic trafficking of cell polarity and cell junction proteins [10-12]. Other than *Drosophila*, caspase-3 associated non-apoptotic functions have been reported in mammalian red blood cells nuclear extrusion [13], neural development [9], liver regeneration process and skin wound healing [14], showing the conservation of the non-apoptotic role in mammals.

Malpighian tubules (MTs) of *Drosophila*, are primarily responsible for excretion and osmoregulation, apart from their role in immune response [15, 16]. They are comprised of two pairs of single layered epithelial tubes, which lodge between the midgut-hindgut junction during embryogenesis and are spatially divided into six segments, the initial, the transitional, the main, lower tubule, upper ureter and lower ureter, reflecting functional differences between the segments. The two physiologically distinctive cell types of invariant number, are, Type I, principal cells (PCs), which are ectodermally derived and Type II, stellate cells (SCs) which are the mesodermal cells, that intercalate into the elongating tubules at stage 13 of embryogenesis [17], among many other cell types that constitute the MTs. Nothing much changes from larva to adult except the shape of the SCs that is remodeled form ‘cuboidal’ in the larval stage to ‘star shape’ in the main segment and ‘bar shaped’ in the initial and transitional segment during pupation [18].

Tubule epithelium produces primary urine in the distal portion via active transport of ions, organic solutes and water from the hemolymph. The PCs are enriched with V-ATPase, along with K^+^/H^+^ exchanger [19, 29] and K^+^ channels [21] in the apical microvilli, and Na^+^/K^+^ ATPase [22], and the Na^+^/K^+^/2Cl^−^ cotransporter [23] on the basal membrane to facilitate the movement of potassium across the PCs. The ensuing imbalance of potassium is counterbalanced by the movement of chloride through chloride channels in the SCs, which results in net movement of water from the hemolymph, through aquaporin DRIP in SCs as well as through paracellular routes [24, 25]. Homeostasis is accomplished by modification of primary urine as it passes down more proximal regions of the tubule before emptying into the hindgut via the ureters [26].

A thought provoking aspect of MTs is their ability to evade apoptosis, unlike most of the larval tissues, during metamorphosis despite the presence of apoptotic proteins in these organs, making it all the more intriguing [27]. The survival has been attributed to the differential gene expression of ecdysone induced genes in MTs compared with salivary glands, which undergo apoptosis [28]. Here we provide evidence to show that caspase-3 homologue, Drice, is involved in a non-lethal role in the development as well as the function of MTs. Drice mutants show grossly misshapened MTs, with imprecisely arranged PCs and SCs, as well as drastic reduction in the number of SCs. The repercussion of Drice deficiency is observed in actin organisation as well as Rho1 expression and localisation of polarity proteins and it also appears to be an important regulator of the acquisition of star shape of the SCs. Aberrant expression of ion channels is observed in Drice mutants, thus affecting the fluid secretion of the tubules. We will be discussing these factors that associate with caspases for the fine tuning of MT development and function. The overall results confirm the non-apoptotic roles of Drice in MTs development, adding to the growing number of studies demonstrating that caspase activation does not always lead to cell death, but rather may promote a variety of non-lethal cellular processes.

## Methods

### Fly stocks

The following mutant alleles were used in this paper: *Drice*Δ*2c8* (Kind gift from Dr. Masayuki Miura, Department of Genetics, The University of Tokyo), *Drice*Δ*1* and *DCP-1(Prev1)* (Kind gift from Dr. Andreas Bergmann, University of Massachusetts Medical School). Principal specific Gal4 driver, c42; stellate cell specific Gal4 driver, c649 (a kind gift from Dr. J.A.T Dow, Institute of Biomedical Sciences, University of Glasgow, UK), Caspase Tracker (DQVD) a *mCD8-DIAP1-GAL4* driven by ubiquitin promoter (kind gift from Dr. Ho Lam Tang, The Johns Hopkins University School of Medicine, USA). UAS responders used: UAS-p35 (Bloomington Stock Centre), G-TRACE-*UAS-RFP; UAS-FLP; Ubi>Stop>GFP-nls* (Bloomington Stock Centre). Appropriate crosses were set to generate UAS-p35; c42Gal4, UAS-p35; c649Gal4 and caspase Tracker/G-trace and the progeny were used for experiments. All stocks and crosses used in this study were maintained on standard *Drosophila* food medium at 24±1 °C.

### Antibodies and immunocytochemistry

MTs from 3^rd^ instar wandering larval stage were dissected in 1X PBS, fixed in 4% paraformaldehyde for 20 min at room temperature, rinsed in 0.1% PBST (1XPBS,0.1%TritonX-100), blocked in blocking solution (0.1% TritonX-100, 0.1%BSA, 10%FCS, 0.1% deoxycholate, 0.02% thiomersol) for 2h at room temperature. Tissues were incubated in primary antibody at 4 °C overnight. After three 0.1% PBST (20 min each) washings, tissues were blocked for 2h and incubated in the secondary antibody. Tissues were rinsed in 0.1% PBST and counterstained with DAPI (1mg/ml, Molecular Probe) for 15 min or phalloidin-TRITC (Sigma-Aldrich, India) at 1:200 dilutions for 2 h at room temperature. Washing was done again in 0.1%PBST and mounted in antifadant, DABCO (Sigma). Primary antibodies used were anti-Dlg (1:20, DSHB), anti-Rho1 (1:20, DSHB), Na^+^/K^+^ ATPase (1:20, DSHB), anti-α tubulin (1:100, Sigma) and Drip (1:1000, Kind gift from Dr. J.A.T Dow).

All preparations were analyzed under Zeiss LSM510MetaConfocal microscope and images were arranged and labeled using Adobe Photoshop7.

### Uric acid deposition

The diameter of the MTs lumen was calculated using profile tool with Zeiss LSM510 MetaConfocal microscope software. The functional analysis of MTs was done by detecting the deposition of uric acid crystals under polarized microscope. The uric acid crystals show birefringence, which is an optical property of any material with a refractive index due to which they appear bright under plane polarized light propagated from a particular direction.

### Scanning Electron Microscopy

MTs were dissected and fixed in Karnovsky’s fixative for 40 minutes on ice, further tissue was post fixed in 1% Osmium tetraoxide in 0.1M PB, pH 7.4 for 1 hour on ice followed by three wash with 0.1 M PB, pH 7.4, 15 min each at 4°C. Samples were immediately dehydrated in a graded series of ethanol (50%, 70%, 80%, 90% and 100%) and then air dried. The dried samples were mounted on carbon taped SEM stubs and sputter coated with platinum. The SEM images were taken using a Carl Zeiss, EVO-18, scanning electron microscope.

## Results

### 1. Compromised caspase-3 activity induces deformities in Malpighian tubule phenotype

We have previously reported the presence of apoptotic proteins in the Malpighian tubules [27] and the developmental and functional defects that ensue due to over expression of apoptotic proteins [29], suggesting that they are important regulators in the MTs, though their exact roles were not deciphered. To delve deep into the understanding of the apoptotic cascade in the Malpighian tubules we examined the phenotype arising out of effector caspase-3, Drice, deficiency. The presence of active caspases in MTs at different stages of development was first confirmed using caspase tracker [30] reporter activity (Supplemental Fig1), and after being convinced of the presence of caspase-3, we went ahead with our studies. To address the question of role of Drice in MTs development, Drice deletion mutants, *Drice*Δ*2c8/Tm6B* and *Drice*(Δ1)/*TM6B*, were chosen; being homozygous lethal at pupal stage, we used homozygous non-tubby larvae for our studies. Taking advantage of the well-defined morphology of wild type MTs, which are thin, elongated, having uniform diameter throughout the length in larval as well as adult stages (Fig1A), we compared them with the mutant tubules. In Drice mutants, *Drice*(Δ1)/*Drice*(Δ1) (Fig1B, C) and *DriceΔ2c8/DriceΔ2c8* (Fig1 D, E, F), there was a distinct loss of tubule morphology and they exhibited gross phenotypes ranging from misshapen tubules to collapsed or sometimes bloated bag (arrows) like structures. Some tubules were twisted throughout the length and were easily broken when straightened manually (Fig1 B, D and F, arrowheads). There was a visible loss of uniform diameter in almost all of the mutant tubules. Statistical analysis of the presence of abnormal tubules in the mutants compared to wild type was significant (data not shown). Since MTs consists of anterior and posterior tubules, we analysed if there was a bias in the defective tubules. Though in most larvae, both the pairs of tubules were malformed, however in some only one pair, out of the 2 pairs was affected. We did statistical analysis and observed that significantly high number of larvae had both pairs of the tubules defective, which was 88% times in comparison to 12% showing defects only in one pair of tubules, which was invariantly the anterior pair (Fig 1G).

**Figure 1:**
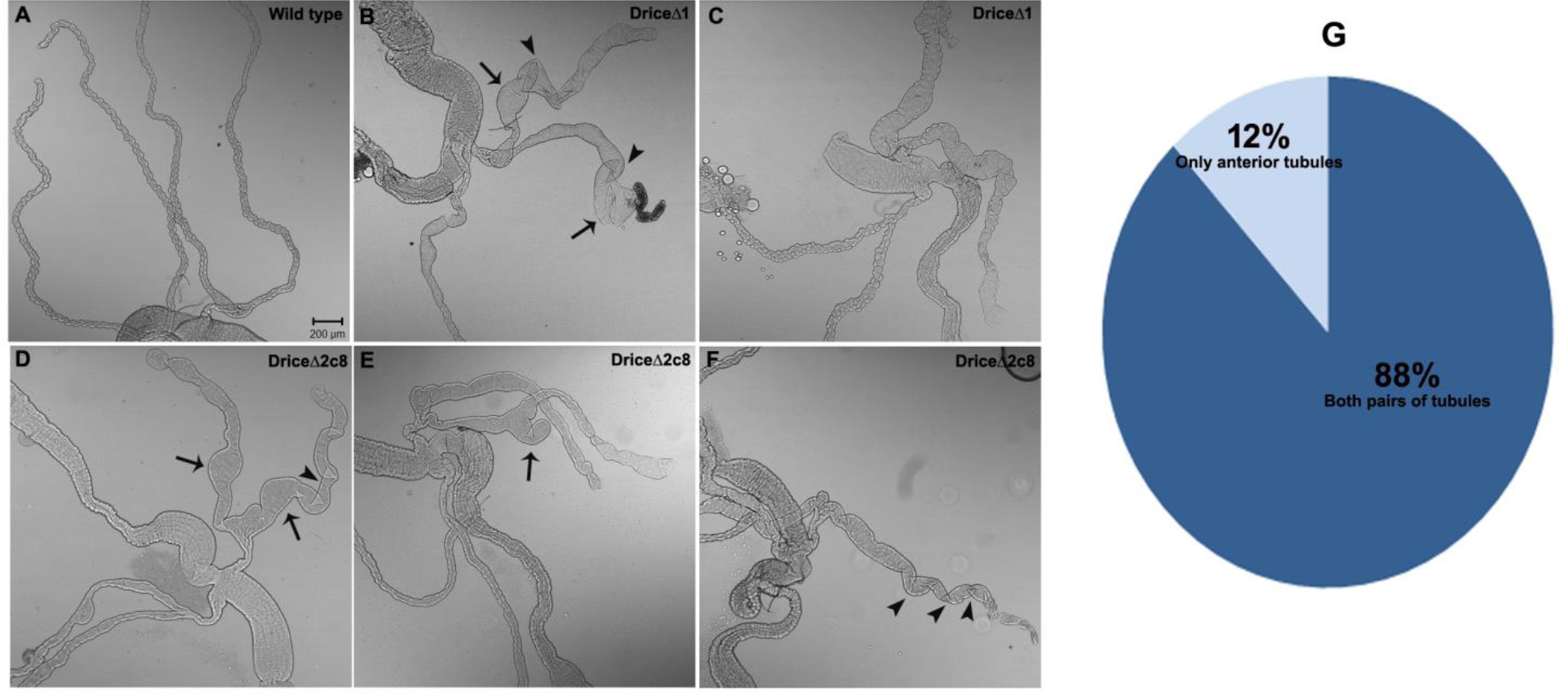
Drice is necessary for proper Malpighian tubule formation. Differential interference contrast images of 3^rd^ instar wild type MTs (A) showing elongated tubules with uniform diameter. *DriceΔ1 (Δ1)* (B, C) and *Drice*Δ*2c8* (D, E, F) mutants displayed failure of tubules elongation and presence of cysts like structures (Arrows) and twists (Arrowheads), scale bar, 200μm. (G) Proportion of all the four tubules affected in comparison to either the anterior or posteriors tubules in *Drice*Δ*2c8* (n=50).

Thorough analysis of several mutant tubules by Scanning Electron Microscope (SEM) showed heterogeneous phenotypes of the tubules ranging from thick, flat and short tubules with irregularly distributed enlargements and constrictions. The smooth basal membrane in wild type (Fig 2A, D), was missing from the mutants which showed distinct disruptions giving the impression of holes in the mutants. Appearance of crevices on the membrane confirmed the loss of tubule formation (Fig 2B, E). Apart from Drice, the second effector caspase is DCP-1 in *Drosophila*, so we examined the MTs in double mutants, *DCP-1(Prev1); Drice*(Δ1), which knocks both the effector casapses, postulating that we should see worsening of the phenotype and as expected, we observed more degenerate and, contorted phenotype of tubules with distinct protuberances (Fig 2 C, F). However, DCP-1 alone did not impair the tubules to the same extent as Drice, suggesting that DCP-1 works in conjunction with DRICE, as reported earlier [31, 32]. These observations showed beyond doubt that caspases are necessary for the morphology and maintenance of Malpighian tubules. Since both the deletion lines showed similar phenotypes, the rest of the experiments were confined to using *Drice*Δ*2c8/Tm6B*, and homozygous larvae are henceforth referred to as *Drice*Δ*2c8*.

**Figure 2:**
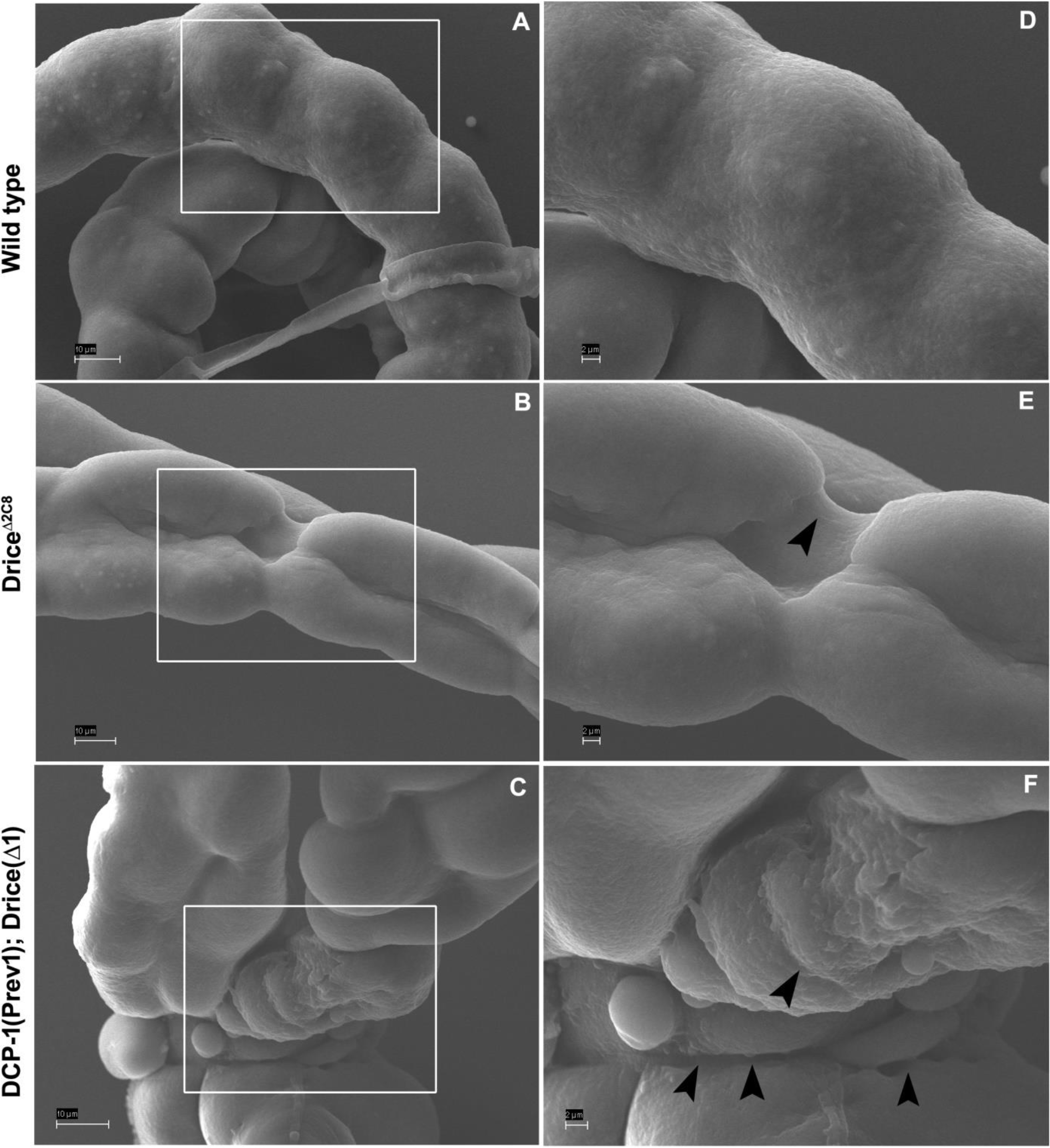
Scanning electron microscopic (SEM) images of Malpighian tubules. The circumference of wild type tubules appeared smooth and the surface was even (A). *Drice*Δ*2c8* had contorted, twisted and irregular appearance with cracks on the surface (B), the notched appearance and the irregularities become more prominent in double mutant *DCP-1(Prev1)*; *Drice*(Δ1) (C). D, E and F are magnified images of the squared boxes in A, B and C. The arrowheads point to perforations on the membrane. Scale bar in A, B, C −10μ and in D, E, F-2μm.

### 2. Tubule length, cell arrangement and number affected following loss of Drice

By the end of embryogenesis a specific number of PCs, fortified with requisite number of SCs make up the tubule tissue, which remains unchanged throughout the life of the fly. The fully developed Malpighian tubules have two cells encircling the lumen, and a patterned arrangement of PCs and SCs, achieved by organised convergent-extension (Fig 3). Since the size of the mutant tubules appeared short, we first established if this observations were significant and measured the length of anterior and posterior the tubules (Fig 4B, D) and compared to wild type (Fig 4A, C). Statistical analysis revealed that the length of the anterior as well as posterior tubules of *Drice*Δ*2c8* mutants were significantly shorter than the wild type (Fig 4E). In order to check whether the disrupted morphology of tubule, also disrupts the cellular arrangement, we next examined the mutant tubules for these readouts. Normal cell arrangement (Fig 4F) shows PCs and SCs intercalated at regular interval, whereas in *Drice*Δ*2c8*, the typical arrangement was completely obliterated and instead of the expected two cells (Fig 3E) many cells were seen clustered along the lumen (Fig 4G), reminiscent of the tubule morphology prior to convergent extension (Fig 3D). As MTs have an almost non-variant number of cells, we next counted the number of PCs and SCs in *Drice*Δ*2c8* mutants (Fig 4H) and observed that there was an overall reduction in the number cells in mutants compared to control. Major effect of Drice deficiency was however, on the number of SCs, where compared to 33 in anterior, and 22 in posterior tubules in wild type, were only 14 and 10 in anterior and posterior tubules respectively in *Drice*Δ*2c8*. Number of principal cells, though not less, was however, variable in mutants. These observations suggested that intercalation of SC is severely affected in mutants. Taken together, these results suggest that there are far reaching consequences of the absence of Drice in MTs.

**Figure 3:**
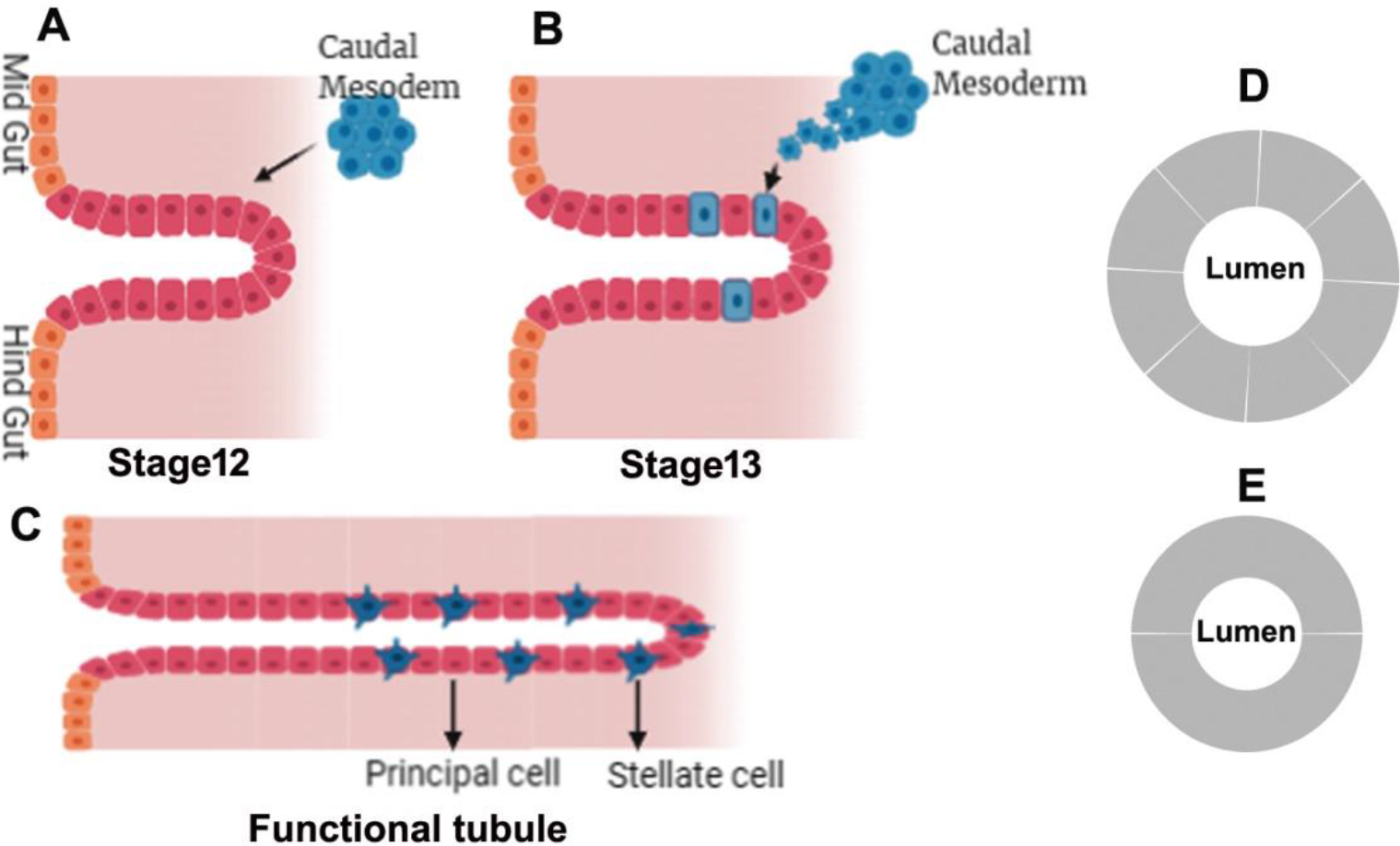
Embryonic development of Malpighian tubules from Stage 12 onwards. After emerging as small bud at the junction of mid gut and hind gut at stage 11, tubule enlarges by synchronous cell division until stage 12 and the caudal mesoderm migrates towards the tubule (A). By stage 13 (B) cell division is complete and further development proceeds by extensive cell intercalation and rearrangement. Stellate cells integrate into the tubule epithelium and undergoes mesenchymal to epithelial transition (B). At this stage, the tubule is short and fat, with six to ten cells encircling the lumen (D). With increase in length and regular arrangement of stellate cells along with decrease in lumen size (C), the final structure of Malpighian tubules is achieved at the end of embryogenesis. At this stage lumen is surrounded by only two cells (E).

**Figure 4:**
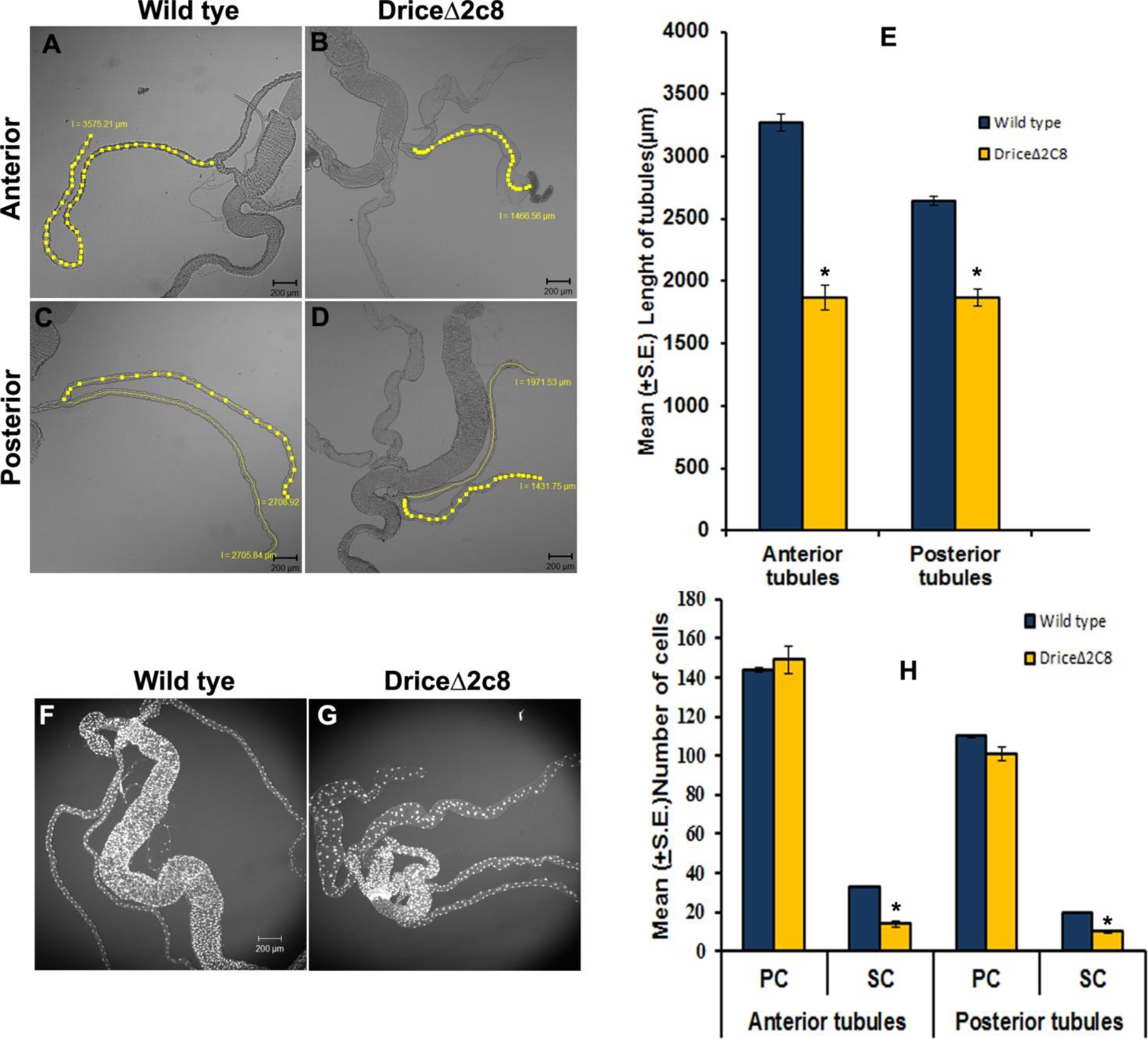
Drice inhibition affects tubule length and SCs number in MTs along with the loss of cellular arrangement. Differential interference contrast image showing measurement of tubule length of wild type (A, C) and *Drice*Δ*2c8* (B, D), Scale bar, 200μm. (E) Bar diagram showing significant differences in the mean length (S.E±) of anterior and posterior tubules between wild type and mutants (n=25 pairs of MTs of each genotype). (F) Wild type tubules have typical cell arrangement with just two cells around the lumen. This arrangement is lost in *Drice*Δ*2c8* (G) and numerous cells present around the lumen. Nuclei are stained by DAPI (Pseudo colour), scale bar, 200μm. Images F and G are projections of optical sections obtained by confocal microscope. (H) Bar diagram showing mean (S.E±) number of PCs and SCs in *Drice*Δ*2c8* compared to wild type (n=25 pairs of MTs of each genotype). The SCs number is more affected compared to PCs in *Drice*Δ*2c8*; statistical analysis was done using t test, * indicates p≤0.05.

### 3. Loss of Drice activity alters the shape of the larval stellate cells

The most conspicuous difference was observed in the shape of the SCs, which are typically cuboidal shaped at larval stage (Fig 5A, B) and transform to star/stellate shape from pharate stage onwards and continue to exist in the same shape in the adult stage (Fig 5C, D). Immunostaining with stellate specific antibody revealed that in contrast to the cuboidal shape of the SCs at 3rd instar larval stage, these cells acquire stellate shape like structure resembling the adult stage, in 3rd instar larval stage itself in *Drice*Δ*2c8* mutants (Fig 5E, F). The formation of filopodia like structures appears at this stage, which could be a prerequisite to the formation of star shape. Similar observations, with more accentuated star shape, were recorded in *DCP-1(Prev1)*; *Drice*(Δ1) double mutant also, (Fig 5G, H).

**Figure 5:**
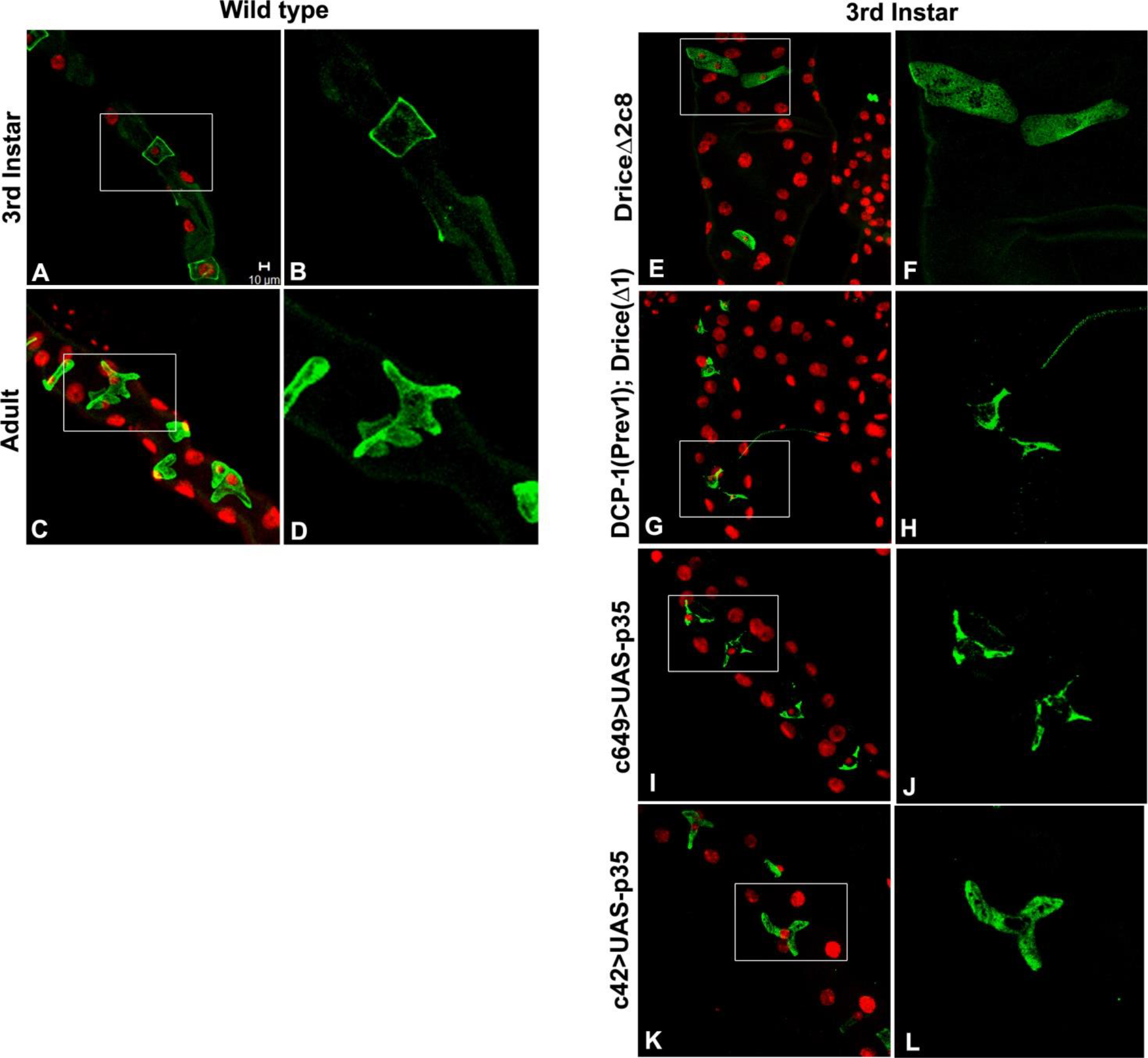
Stellate cell shape is affected by Drice inhibition. SCs are identified by the expression of Drip, in wild type MTs they have a cuboidal shape at 3^rd^ instar larval stage (A, B) and star shape at adult stage (C, D); while in absence of Drice, in *Drice*Δ*2c8*, these cells lose their shape and acquire elongated shape (E and F), these features are more accentuated in double mutant, *DCP-1(Prev1)*; *Drice*(Δ1) (G and H) with filopodia like extensions. Down regulation of Drice, specifically in SCs, c649>UAS-p35 (I and J), as well as down regulation in PCs (K and L) also affect the SCs shape in a similar fashion. Image B, D, F, H, J and L are magnified image of A, C, E, G, I and K respectively. Nuclei are stained by DAPI (Pseudo color red). All images are projections of optical sections obtained by confocal microscope, scale bar, 10μm.

To confirm whether the transformation in the SCs shape was indeed due to reduced Drice, we used UAS p35 to specifically down regulate Drice in SCs using c649 Gal4 and observed similar change in the morphology of the SCs converting them into star shaped in third instar larval stage (Fig 5I, J). Further we also checked the effect of inhibiting Drice in PCs using c42 Gal4 and observed that inhibition of Drice in PCs also affected the SCs shape in a similar manner (Fig 5K, L). These observations suggested confirmed the role of Drice in transformation of the shape of SCs from cuboidal to star shape and an inter-cellular dependence between the two cell types in MTs for regulation of cell shape. The emergence of the star shape of SCs in larval stage itself in contrast to the regular cuboidal shape, suggested that effector caspases restrains the attainment of stellate shape until adult.

### 4. Disorganised cytoskeleton in Drice mutants leads to convergent extension defects

The lack of visible proximo-distal (P-D) elongation in Drice mutants can be attributed to defective cytoskeleton organization, which is essential for the formation and maintenance of the tubule morphology [33]. Hence we sought to examine the actin and tubulin organisation in 3^rd^ instar larvae. F-actin was observed by staining with phalloidin and was found to be localized in cell cortex (Fig 6A) and particularly at apical surface (luminal side) of all tubule cells (Fig 6B) in wild type, which is consistent with earlier report [33]. In *Drice*Δ*2c8* mutants and double mutants, however there was substantial increase in the expression of F-actin with complete disruption of its organization (Fig 6B, C and D). The actin filaments appeared very dense and more diffused throughout the tubules. The disarrangement of the actin was more pronounced in the *DCP-1(Prev1)*; *Drice*(Δ1), double mutants (Fig 6C, D). The presence of actin at the lumen (Fig6 F-H) was although reminiscent of the wild type, but the expression in mutants was proportionately very high. Bundles of actin fiber around the SCs (Arrowheads, B, D) were more compact and pronounced in Drice mutants, than wild type. Discontinuous localization of actin along the length indicates that the actin network produces local constrictions as well as expansion along the circumference of the tubule morphology, consequently giving rise to irregularity in shape of tubules which was clearly visible in the SEM images of the tubules (Fig 2). These data indicate that Drice is required for normal spatial distribution of actin in MTs during development.

**Figure 6:**
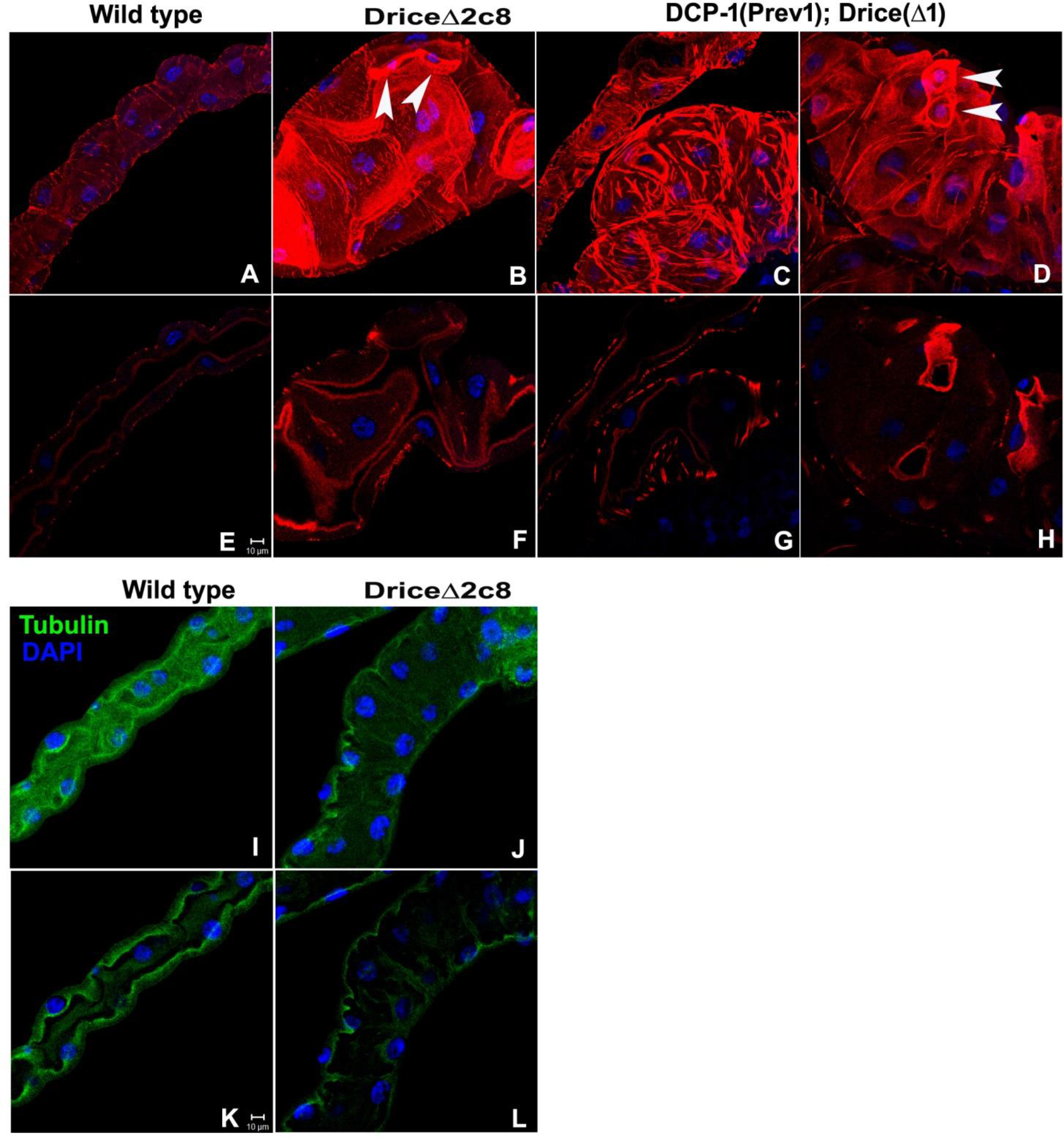
Disruption of cytoskeletal elements in Drice mutants. F-actin organization on cell membrane, in cytoplasm, and at the lumen of MTs of wild-type third instar larvae (A and E) is seen to be disrupted in *Drice*Δ*2c8* (F and F) and *DCP-1(Prev1)*; *Drice*(Δ1) (C, D, G and F). E, F, G and H are section images of A, B, C and D respectively, showing lumen arrangement in tubules. Images A-D are projections of optical sections obtained by confocal microscope, scale bar 10μm. Arrowheads show enhanced actin accumulation in SCs of *DCP-1(Prev1)*; *Drice*(Δ1). Wild type tubules showed apical α-tubulin expression (I and K), which is seen to be disrupted in *Drice*Δ*2c8* (J and L). Nuclei are stained by DAPI (blue). Images I-J are projections of optical sections obtained by confocal microscope, images K and L are single section of J and L respectively, scale bar 10μm.

Expression of tubulin was observed by staining with anti-α tubulin antibody and unlike actin, the expression of tubulin was significantly less in Drice mutants as compared to wild type. Further it was observed that there was distinct localisation of tubulin on the lumenal region of wild type (Fig 6I, K), which was missing from the lumenal region and appeared to be more on the basal membrane in the mutant (Fig 6 J, L).

We next checked the integrity of apico-basal polarity of MTs cells, by observing the localization of Disc large (Dlg), a basolateral membrane protein. The normal wild type pattern of Dlg (Fig 7A, E) was absent in the mutants and the distribution of Dlg was severely affected. Dlg could be seen forming a thick and diffused line in mutant (Fig 7B, F) unlike that of wildtype where it is strictly on the membrane. Moreover in double mutant, Dlg localization was completely disrupted (Fig 7C, G). These results indicate that probably the endocytic trafficking of Dlg is disturbed, reminiscent of the function of Drice in endocytic trafficking during *Drosophila* tracheal formation [12]. Moreover, loosening of cell contacts and multilayered formation, gaps between tubular cells and rounding up of cells as protuberances (Fig 2), which could be due to improper cell-cell junctions, was observed in Drice inhibited condition.

**Figure 7:**
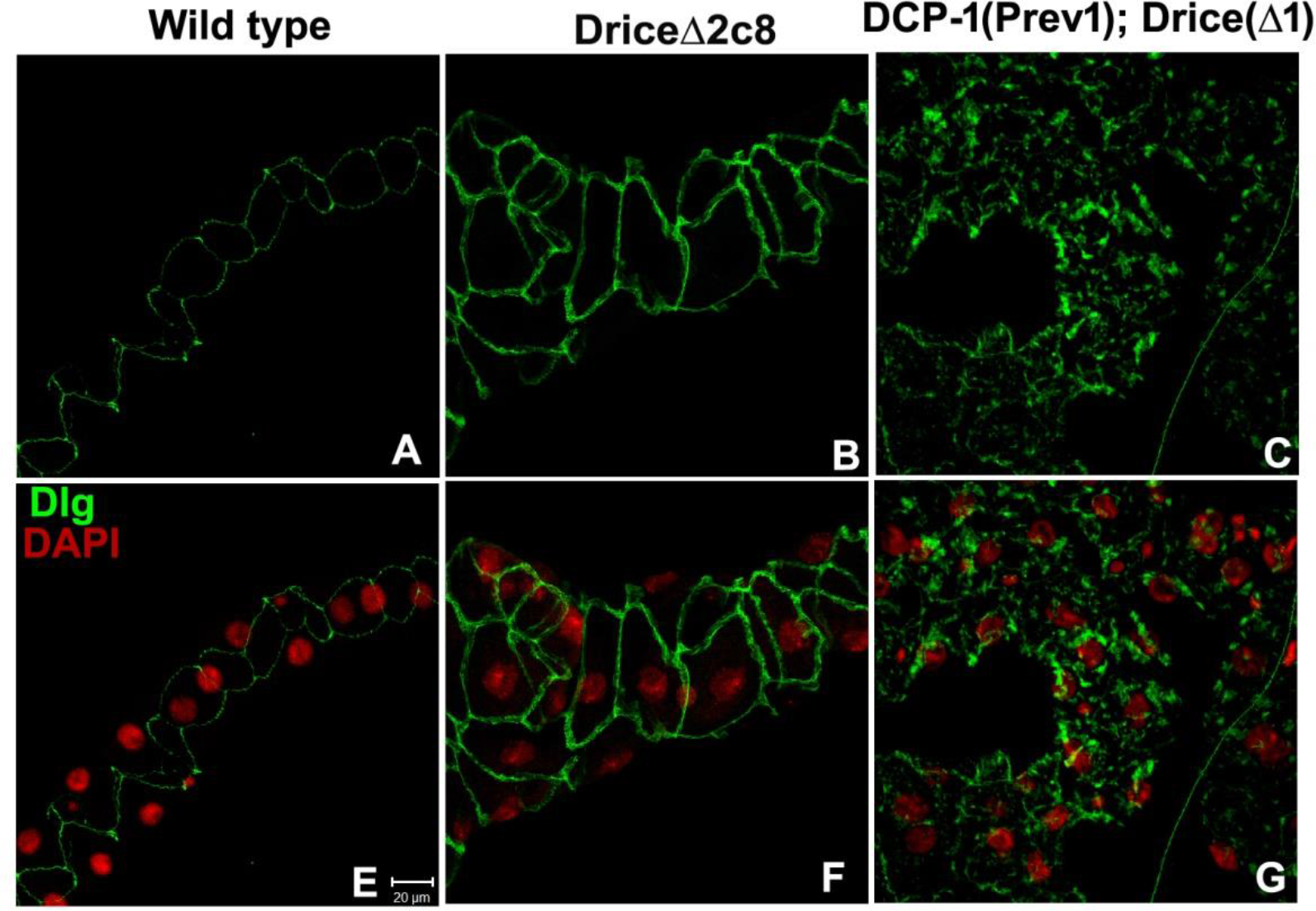
Inhibition of Drice results in disruption in Disclarge (Dlg) localization. Dlg is localized to the baso-lateral lining of MTs in wild-type third instar larvae (A and E), in *Drice*Δ*2c8* (B and F) and *DCP-1(Prev1)*; *Drice*(Δ1) (C and D) Dlg localization is disrupted. Nuclei are stained by DAPI (Pseudo colour red) in E, F, G. All images are projections of optical sections obtained by confocal microscope, scale bar, 20μm.

### 5. Disruption of Rho1 expression and localisation in absence of Drice

Extensive disruption of cell arrangement, cell shape and actin organisation suggests involvement of cytoskeletal dynamics. During development, small RhoGTPases control the precise cell shape changes and movements that underlie morphogenesis. RhoGAP, crossveinless-c (cv-c), is known to regulate spatial organization of F-actin in tubule cells via inactivating small GTPase, Rho1 [33]. In *cv-c* mutants, F-actin network is disrupted and diffused, because of which MTs collapse to form cyst like sack structures. Thus similar collapse in tubular morphology, along with actin disorganization in absence of caspase-3 in tubules encouraged us to examine the involvement of Rho1 in this process. Immunostaining for Rho1 revealed that the expression in *Drice*Δ*2c8* (Fig 8B) and *DCP-1(Prev1);DriceΔ1*, double mutant (Fig 8C), was considerably high compared to wild type (Fig 8A), indirectly suggesting a compromised cv-c activity. Transverse section images showed that Rho1 localisation is more intense on the membrane (Fig8 B’ C’) unlike the wild type (Fig 8A’). These results indicate that caspase-3 is probably regulating tubules morphogenesis via Rho1 either directly or through cv-c.

**Figure 8:**
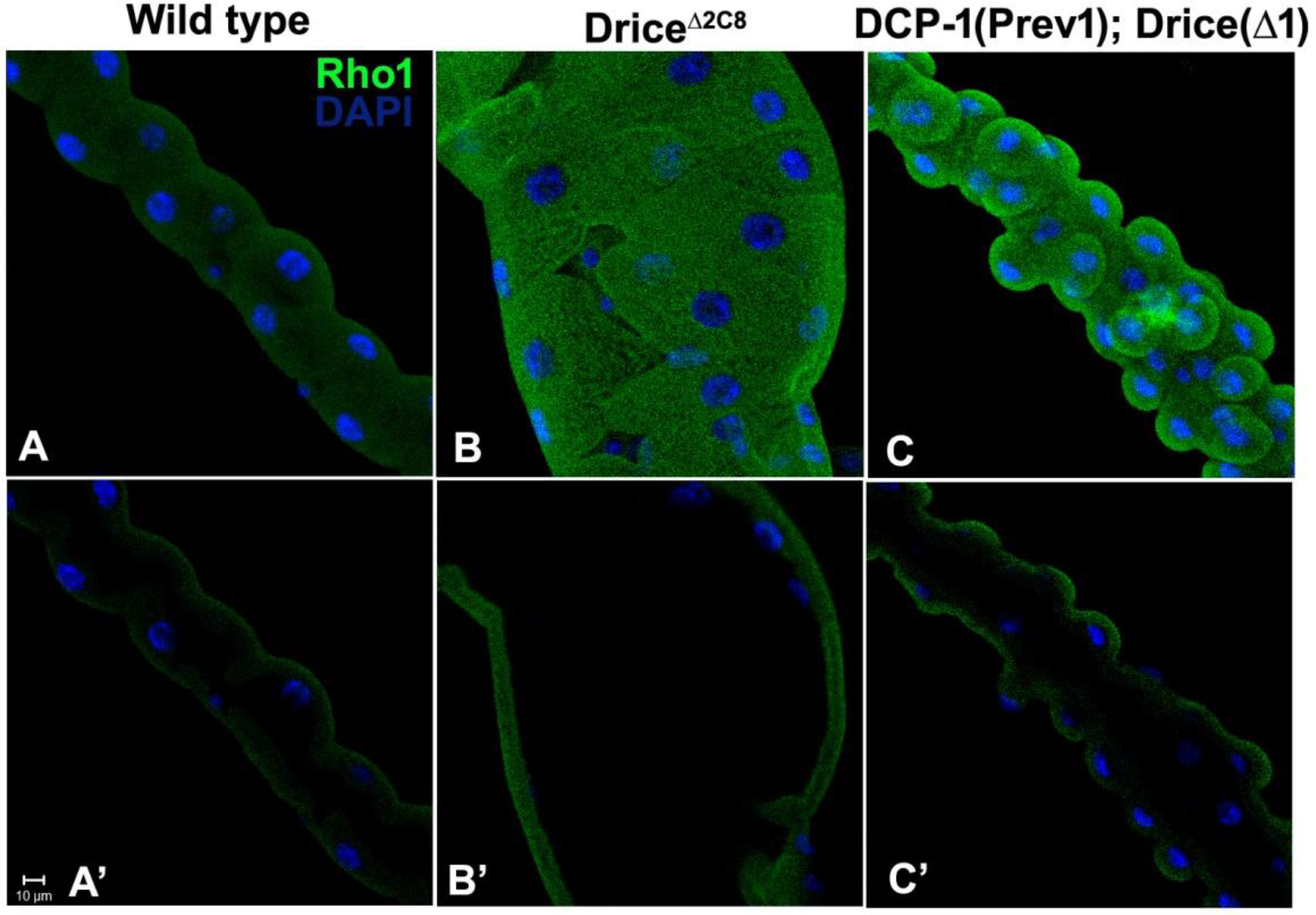
Rho-1 expression and localisation is affected in the absence of Drice. Minimal expression of Rho1 is seen in wild type tubules (A, A’), while in *Drice*Δ*2c8* (B, B’) and *DCP-1(Prev1)*; *Drice*(Δ1) (C, C’) Rho1 expression is increases. Nuclei are stained by DAPI (blue). A, b and C are images are projections of optical sections obtained by confocal microscope while A’, B’ and C’ are single sections, scale bar, 10μm.

### 6. Renal function get affected in the absence caspases

Precise cellular arrangement is a prerequisite for absolute function of any organ. Since gross morphological defects were observed in Drice mutants we assessed the corollary on the function of the tubules. The first difference was observed in the lumen of mutant tubules, which has multiple defects, was either not formed properly, or there was no lumen at all or the lumen size was greatly enlarged (Fig 9B) compared to wild type (Fig 9A). In wild type maximum width of lumen was in a range of 54μm while in Drice mutants it was in a range of 168 μm, which, needless to say was significantly high and likewise diameter of the narrowest region in wild type tubule was also significantly less in comparison to narrowest region of mutant tubules (Fig 9G). A single tubule showed variable phenotypes where in some regions lumen was not formed, whereas in the adjoining region it was very much enlarged. A prominent feature on the inside of the lumen were the infoldings of the apical membrane, at regular intervals, in the proximal direction (Fig 9C, arrows), probably a mechanism for unidirectional flow of urine. These were however not sculpted in Drice mutants, as well as a noticeable reduction in the thickness of the membrane was also observed. These defects could be consequential to abnormal actin and tubulin organization, resulting in loss of typical lumen formation in the mutants. The functional competence of tubules was adjudged by observing the uric acid crystals under polarized light, as these crystals exhibit birefringence. The mutant tubules showed enormously high accumulation of uric acid crystals (Fig 10B) in comparison to WT tubules (Fig 10A), indicating a complete breakdown of the physiological functions. The prominent enlargements in Drice mutant MTs appeared fluid filled with dark crystalline particles floating which were never observed in wild type (Fig 10 C, D, E and F). It is possible that the accumulations of dark crystalline structures and uric acid crystals in mutant tubules, is due to blocked lumen.

**Figure 9:**
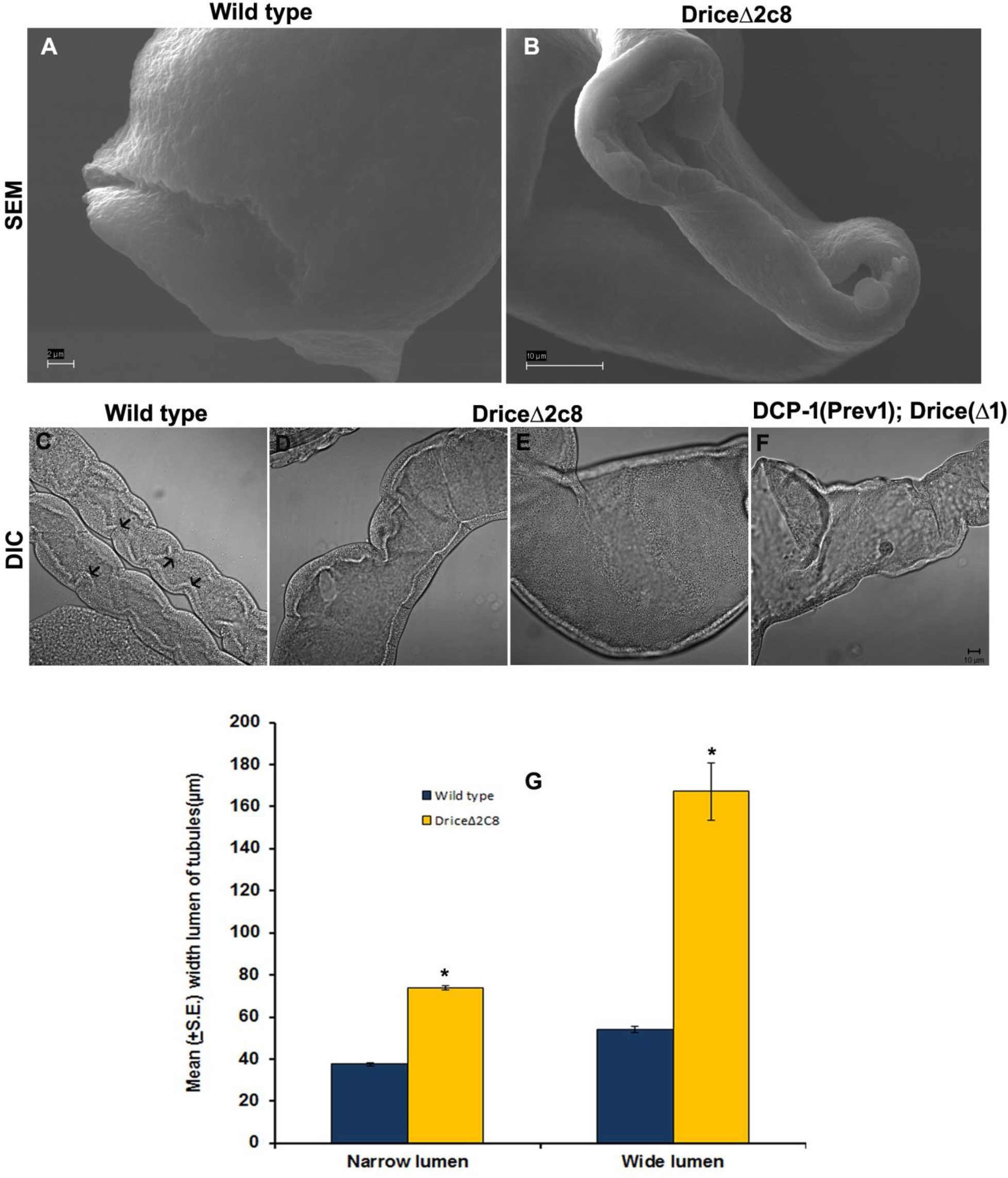
Typical structure of lumen and its size was severely affected by caspase inhibition. Scanning electron microscopic images of Malpighian tubules showing lumen architecture in wild type (A) and *Drice*Δ*2c8* (B). Image A is at higher magnification (scale bar 2 μm), while (B) is at lower magnification (scale bar 10μm). These images show that lumen is extremely enlarged and flattened in mutant tubule. DIC images showing lumen of wild type (C) with infoldings which are shown by arrows, while *Drice*Δ*2c8* (D, E) and *DCP-1(Prev1)*; *Drice*(Δ1) (F) do not show such infoldings of luminal wall, scale bar is 10μm. (G) is a bar diagram showing mean (S.E±) lumen width of Malpighian tubules in *Drice*Δ*2c8* compared to wild (n=25 pairs of MTs of each genotype), lumen was categorized into wide and narrow lumen type. Statistical analysis was done using t test,* indicates p≤0.05.

**Figure 10:**
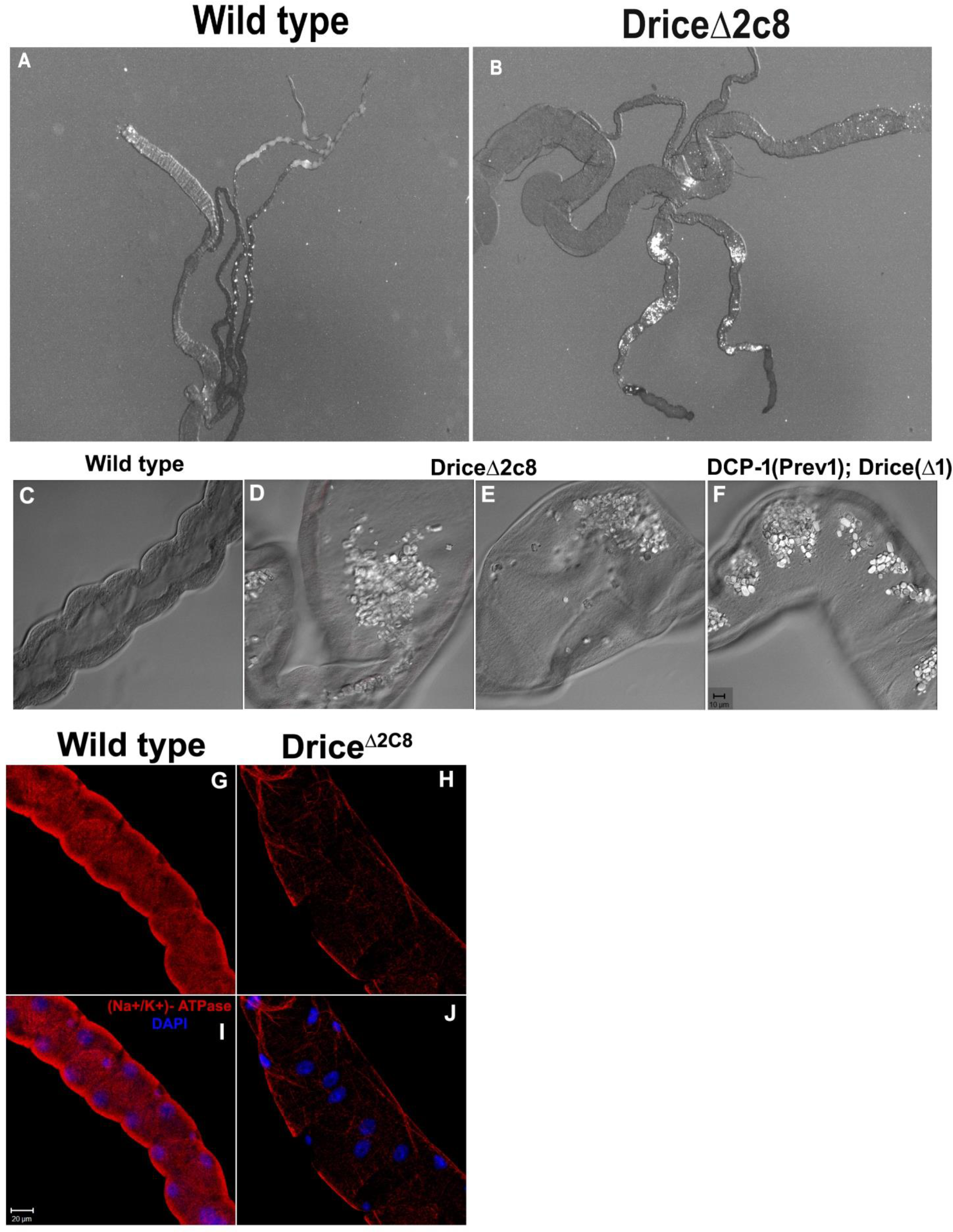
Caspase inhibition results into physiological defects in Malpighian tubules. Deposition of uric acid crystals in polarized light was found to be normal in wild type 3^rd^ instar larvae (A) whereas *Drice*Δ*2c8* 3^rd^ instar larvae it was extremely enhanced (B). Crystalline depositions were observed in lumen of *Drice*Δ*2c8* (D, E) and *DCP-1(Prev1)*; *Drice*(Δ1) (F) by Differential interference contrast (DIC) imaging, which were completely absent in wild type condition(C), scale bar is 10μm. Caspase is required for proper localization of ion transporter Na^+^/K^+^ ATPase, caspase inhibition causes decreased expression of Na^+^/K^+^ ATPase in *Drice*Δ*2c8* (H and J), in comparison to wild type (G and I). Images G-J are projections of optical sections obtained by confocal microscope, scale bar, 20μm.

The presence of Na^+^/K^+^ ATPase pump on the basal membrane of PCs has an important role in the production on fluid in the lumen. We compared Na^+^/K^+^ ATPase level in Drice mutants with that of Wild type (Fig 10G and I), and observed that its level was very much reduced in mutants (Fig 10H and J), signifying compromised physiological function in the effector caspase mutants.

## Discussion

In this paper, caspase-3 has been qualified to be of paramount importance for the integrity of MTs of *Drosophila*. The evasion of apoptosis during metamorphosis, despite the presence of apoptotic proteins had intrigued us, and investigation into the rationale for their presence led us to establish that MTs fail to achieve the functional 3-dimensional structure and organisation in the absence of effector caspase-3, Drice. Although MTs normally do not undergo apoptosis, but they are capable of undergoing cell death, by autophagy as well as apoptosis under specific conditions [27, 28], which has also been reported in tracheal Dorsal trunk which requires Drice for development, but at the same time can undergo apoptosis if induced [12]. These facts suggest that the activity of Drice is kept below threshold, required for apoptosis.

During embryonic development, the MTs undergo convergent extension, soon after cell division is complete and appropriate cell number is reached, in the proximo-distal axis to achieve the transformation from a bulbous structure having 10-12 cells surrounding the lumen, to two cells encircling the long thin tubules [18, 34]. The final structure of MTs, is achieved by cell motility that drives intercalation, an organ autonomous process [35, 36]. The convergent extension movements with concomitant change in cell partners and cell shape, reminiscent of germ band extension [37], are dependent on precise remodelling of actin cytoskeleton and zonula adherens junctions to accommodate change in tissue shape [37]. SEM images of mutant MTs clearly show defects in ZA junctions in Drice mutants, which gets more punctuated in Drice-Dcp1 double mutants, and is validated by grossly defective actin organisation as well as tubulin arrangement. Since actin and tubulin are also known substrates of caspase-3 [38, 39], therefore in the absence of Drice activity, they exhibit disrupted distribution with major increase in the cortical actin. These changes affect cell motility that drives intercalation and hampers with the signalling pathways that regulate MT development. The short MTs in the absence of Drice are reminiscent of tracheal dorsal trunk elongation defects in Drice mutants [12].

Junctional proteins are particularly important for tissue remodelling [37, 40], and inappropriate pattern of Dlg in Drice mutants, as well as *DCP-1(Prev1)*; *Drice*(Δ1) double mutant; and other junctional proteins (data not shown) may interfere with correct patterning of cells. The membrane proteins are stabilised by linking to the cytoplasmic scaffolding proteins and actin/spectrin cytoskeleton. The presence of Drice on the membrane [27], could be additionally involved in providing a scaffold to retain them appropriately on the membrane. Absence of Drice resulted in complete redistribution of actin and other membrane proteins, which is also observed in trachea where endocytic trafficking of junctional proteins is regulated by Drice. These defects may consequently lead to the non-typical arrangement of PCs and less number of SCs.

The mature architecture of MTs entails a concerted action of cv-c, encoding RhoGAP, with Rho1 being the primary target in tubules [33]. The present results show activation of Rho1 in Drice inhibited condition, suggesting that Drice could be negatively regulating Rho1 expression and its overexpression, could probably be the reason behind the uncoordinated actin organization. Whether the effect on Rho1 by Drice is direct or through cv-c, has to be established. Moreover it has also been reported that four family members of RhoGTPases i.e, Rho1, Rac1, Rac2 and Cdc42 have conserved caspase cleavage sites and caspases could regulate a cell migratory behaviour by inactivating RhoGTPases [41].

Similar phenotypes are observed in mutants of *zipper*, encoding non-muscle myosin heavy chain (NMHD), and *ribbon* and *raw* needed for the sub cellular localisation of NMHD show similar phenotypes when mutated [42, 43]. An additional mechanism of elongation in tubule could be the formation of lamellipodial protrusion, as observed in Xenopus notochord formation [44]. Polarized basolateral cell motility underlies invagination and convergent extension, and their dependence on extra cellular matrix (ECM) to bring about directional elongation in one axis. Since these factors are dependent on the basal membrane [45], which is highly disrupted in Drice mutants, confirmed our belief that the failure of MTs formation could be dependent upon these factors as well. As Drice is a caspase, the first conclusion that can be derived is probably the above mentioned products/proteins are targets of Drice and therefore signaling appears to be disrupted globally, though at present the answer still eludes us.

SCs are mesodermal in origin and they undergo mesenchymal to epithelial transition [17] and intercalates in the elongating tubule during stage 13 as shown in Fig 3. Absence of Drice impinges on the number of SCs as well as shape. Significantly less number of SCs get intercalated which could be due to defects in actin organisation, membrane proteins, as well as defects in the ECM, thus influencing intercalation of SCs during elongation of MTs.

The star shape of SCs, which is a characteristic feature of adults responsible for maximising the surface area of SCs for function, is observed at the larval stage itself in Drice mutants suggesting that caspases has a role in restraining the shape of the SCs, similar to the negative regulation of speed of thorax closure [11]. Teashirt, specifically expresses in the SCs, is required for cell shape changes as well as physiological maturation [18], however, since there is no change in teashirt expression in Drice mutants compared to wild type (Data not shown), it is likely that Drice is acting down stream of Teashirt. This change in shape, we believe is a stage specific requirement and attainment of this shape at the larval stage could result in functional impairment and be catastrophic. Precise development of an organ is closely linked to its functional status. Similar to vertebrate’s nephrons, the MTs are tubular organ and correct arrangement of cells as well as the lumenal architecture, is of paramount importance for the formation of lumen. Defects can lead to ion and water imbalance in the haemolypmh, which can lead to lethality or development of polycystic kidney diseases like phenotype as seen in in humans where nephrons get highly distended [46]. In higher organisms many organs including salivary glands, mammary glands, liver and kidney necessitate unerring formation of epithelial tubes for proper function.

Primary urine is secreted by main segment cells, driven by ion transport across the epithelium. In *Drosophila* two physiologically distinctive cell types: PCs and SCs drive this process. PCs constitute about 80% of all tubule cells, and are responsible for the transport of cations and organic solutes [47]. SCs are scattered around the PCs and are responsible for water and chloride ion (Cl−) flow through aquaporin water channels (Drip) and chloride channels respectively. Therefore it is conceivable that anomalous increase in lumen diameter, compounded with defects in Na^+^/K^+^ ATPases, due to inadequacies of basal membrane; and probable reduction in chloride conductance due to reduced number of SCs, could be ascribed to defective function for generation of primary urine and formation of excessive uric acid crystals in Drice mutants. The reduced number of SCs may also impact water movement, as SCs are acquainted with Drip, the water channels hence reduction in number of cells would lead to reduction in water movement. We hypothesize that a fine balance of active caspase, which is kept below threshold, is essential for development and functional integrity. The results presented in this paper are expected to reflect unequivocal evidence about non-apoptotic role of caspase Drice in regulating cell movement and intercalation during elongation of MTs development. It demonstrates that effector caspase, well known for its role in apoptosis has a role in negatively regulating cell movement during morphogenesis, thus demonstrating a new and unexpected link between apoptosis and cell mobility.

## Acknowledgements

We thank Bloomington Drosophila Stock Center, Dr. J.A.T. Dow, Dr. Masayuki Miura, Dr. Andreas Bergmann and Dr. Ho Lam Tang for sharing their fly stocks. We thank Dr. N.V. Chalapathi Rao, Banaras Hindu University, for Scanning electron microscope facility. We also thank Dr. Hena Firdaus, Central University for Jharkhand, for providing polarized microscope facility. This work was supported by research grant from Department of Science and Technology and Department of Biotechnology Government of India, New Delhi. We also thank Department of Science and Technology, India, for providing National Facility for confocal microscope. Shainy Ojha was supported by a research fellowship from the Council of Scientific and Industrial Research, New Delhi and research grant from SERB, New Delhi is highly acknowledged.

