## Supplementary Figure 1 for "Non-apoptotic function of caspase-3 in morphogenesis of epithelial tubes of *Drosophila* renal system"

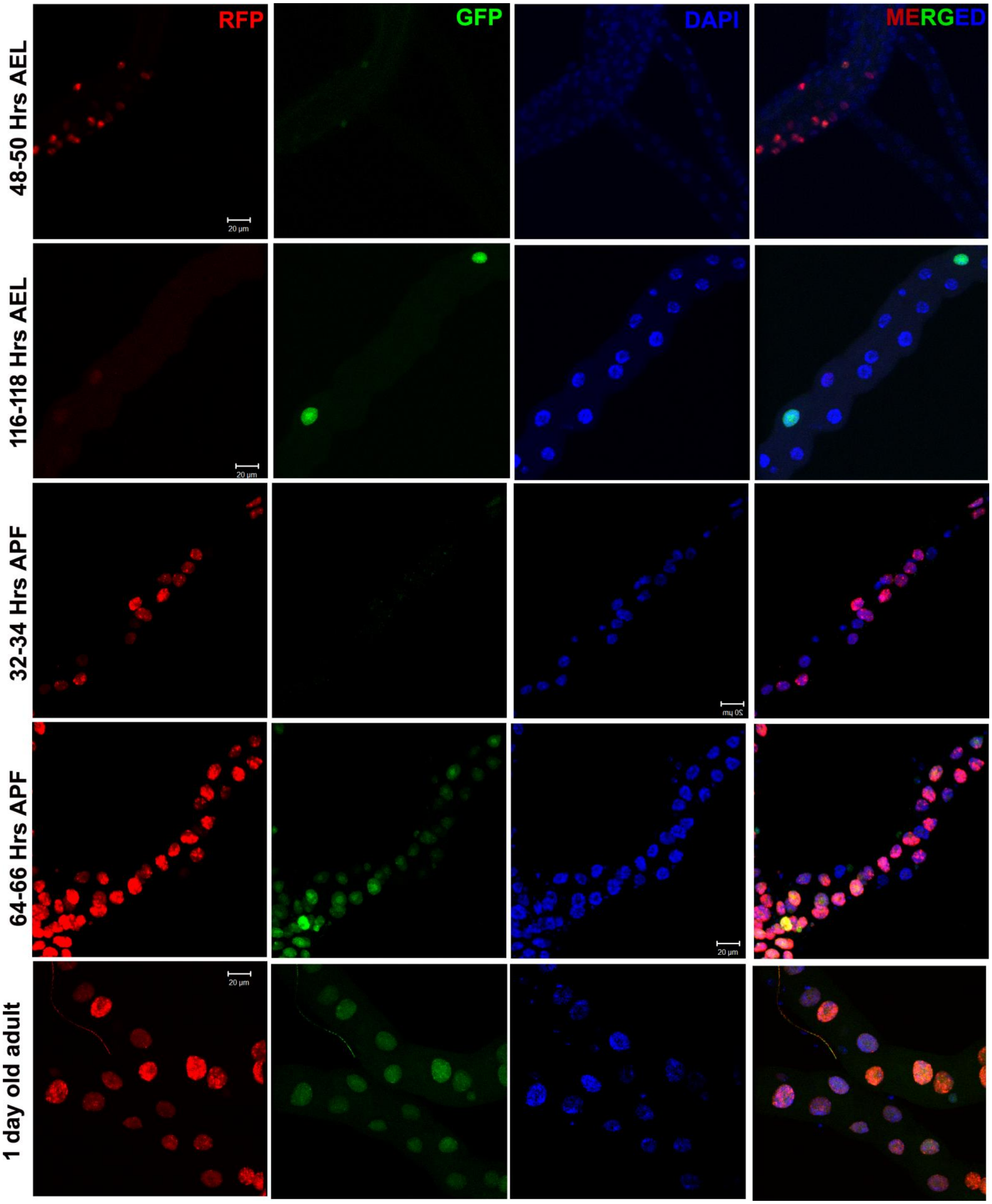

**Supplementary Fig 1: Caspase activity in Malpighian tubules (MTs) at different stages.** Dual colour Caspase Tracker biosensor showing expression of transient caspase activity with red fluorescence RFP, past caspase activity is shown with green fluorescence GFP and chromatin was stained with DAPI in MTs at different stages. The merged panel shows the overlap of past and present caspase activity in the nucleus. All images are projection of optical sections obtained by Zeiss LSM 510 Meta Confocal microscope, scale bar, 20 μm.
